# CoBATCH for high-throughput single-cell epigenomic profiling

**DOI:** 10.1101/590661

**Authors:** Qianhao Wang, Haiqing Xiong, Shanshan Ai, Xianhong Yu, Yaxi Liu, Jiejie Zhang, Aibin He

## Abstract

An efficient, generalizable method for genome-wide mapping of single-cell histone modifications or chromatin-binding proteins is so far lacking. Here we develop CoBATCH, <u>co</u>mbinatorial <u>b</u>arcoding <u>a</u>nd <u>t</u>argeted <u>ch</u>romatin release, for single-cell profiling of genomic distribution of chromatin-binding proteins in cell culture and tissue. Protein A in fusion to Tn5 transposase is enriched through specific antibodies to genomic regions and Tn5 generates indexed chromatin fragments ready for the library preparation and sequencing. Importantly, through a combinatorial barcoding strategy, we are able to measure epigenomic features up to tens of thousands single cells per experiment. CoBATCH produces not only high signal-to-noise features, but also ~10,000 reads per cells, allowing for efficiently deciphering epigenetic heterogeneity of cell populations and subtypes and inferring developmental histories. Thus, obviating specialized device, CoBATCH is easily deployable for any laboratories in life science and medicine.

## MAIN TEXT

Single cell Transcriptomics^1, 2^ and single cell Epigenomics^3–8^ have seen great stride and invaluable utilities in revealing cell fate specification and population heterogeneity during developmental processes and pathological alterations. Development of efficient and robust single-cell ChIP-seq (Chromatin Immunoprecipitation in parallel with Sequencing) methods for profiling epigenetic landscape far lagged behind, despite numerous efforts and progress on low input cells have been made^9–13^. Here, we developed a versatile strategy, dubbed in situ ChIP for a small amount of materials and combinatorial barcoding and targeted chromatin release (CoBATCH) for single-cell level.

For in situ ChIP with low-input cells, we placed the N terminal of Tn5 transposase to Protein A (PAT) for targeted tagmentation and the one-step sequencing library preparation, obviating DNA end repairing and adaptor ligation. Varying cell numbers of mouse embryonic stem cells (ESCs) were permeabilized and incubated with antibodies (H3K27ac, H3K27me3 and H3K4me3) before addition of PAT to tether Tn5 transposase to the specific histone marks or proteins of interest through protein A (Figure 1A). Targeted tagmentation and indexed fragment release were performed, followed by PCR enrichment for the sequencing ready library in one tube without need of DNA extraction and purification. We acquired high quality in situ ChIP signals for all three histone marks as few as 100 cells, as demonstrated by specific inspection of a few positive peak regions defined by public data from ENCODE (Supplementary Figure 1A-C). Notably, due to the high signal-to-noise advantage of this approach, the sequencing depth of the unique non-duplicated reads of ~5M generated highly reproducible data as demonstrated by peak intersection. Highly enriched H3K27me3 signals, a repressive mark preferentially found in regions of lower chromatin accessibility^14^, confirmed that in situ ChIP method was compatible not only to active histone marks, largely overlapping with open chromatin regions, identified by ATAC-seq, but also to closed chromatin regions. Further global analysis supported the reproducibility and robustness of in situ ChIP in low-input cells (Supplementary Figure 1D-F).

**Figure 1.**
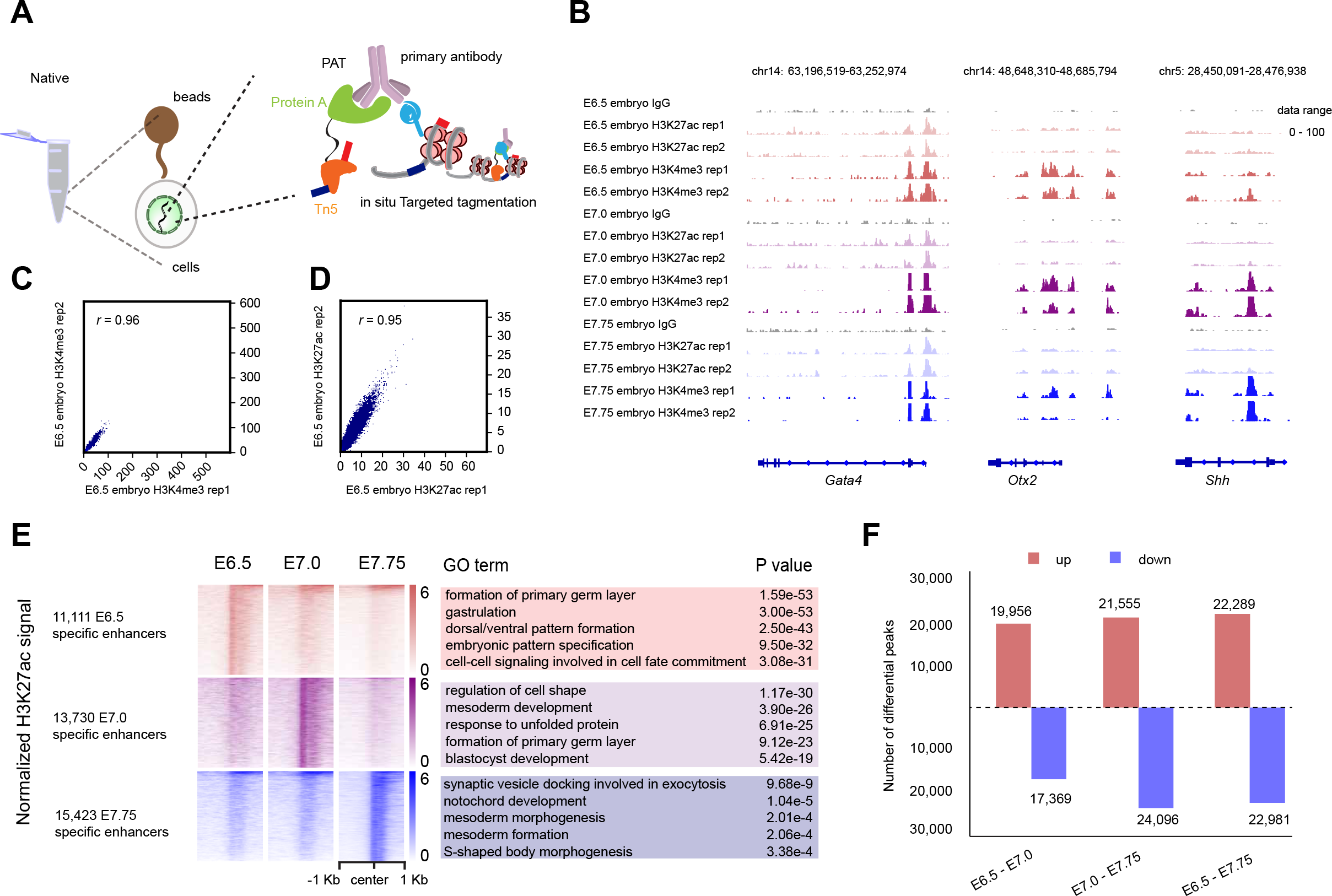
Native in situ ChIP in intact tissues during mouse gastrulation. (A) Illustration of the key steps for low input in situ ChIP. Blue lollipop represents chromatin binding proteins or histone marks, H3K27ac, H3K4me3, and H3K27me3. PAT denotes Protein A and Tn5 fusion protein. (B) Track view showing normalized H3K27ac and H3K4me3 signals with native whole embryos (E6.5, E7.0, and E7.75) at selected loci. (C-D Scatter plot showing correlation between biological replicates. Seen also in Methods. (E) Left: Normalized H3K27ac signals at ± 1 kb of stage-specific enhancer centers is plotted. Stage specific enhancer peaks are identified as below: broad enhancer peaks are called by MACS2 using two merged replicates of each stage excluding regions at the proximal promoter of genes. This was calculated by “mergePeaks” in Deeptools. Stage specific enhancers associated GO terms (Biological Process) were identified by GREAT. (F) Dynamics of H3K27ac peaks during the gastrulation process. Violet bars (up) indicate H3K27ac peaks uniquely present in the later stage and red bars (down) indicate H3K27ac peaks uniquely present in the former stage. “mergePeaks” function was used to calculate unique peak numbers.

Next, we tested whether in situ ChIP method can be directly implemented to intact tissues with scarce cells while elimination of physical isolation of individual cells. To this end, we profiled H3K4me3 and H3K27ac binding in developing mouse embryos to reveal the epigenomic landscape dynamics from the onset of gastrulation from embryonic day 6.5 (E6.5) to the end at E7.75. Track view for specific target regions for H3K4me3 and H3K27ac in situ ChIP using single embryos each experiment showed the similarly high enrichment and extremely lower background in all groups with varying cell numbers from E6.5 (~660 cells), E7.0 (~4,500 cells) and E7.75 (~15,000 cells)^15^ (Figure 1B). Genome wide scatter plots indicated a high reproducibility in single E6.5 embryos (Figure 1C,D). We also validated the true peaks of H3K27ac as an active enhancer marker^16^, called from our method, as demonstrated by over-representation of related Gene Ontology (GO) terms, such as “formation of primary germ layer” and “gastrulation” in E6.5, “mesoderm development” in E7.0 and “mesoderm morphogenesis” and “S-shaped body morphogenesis” in E7.75, coinciding with the stage-specific developmental events (Figure 1E). Our data enabled identification of a large number of dynamic enhancers during gastrulation, providing a resource to understanding of epigenetic mechanisms and also inferring pivotal transcription factors with instructive roles in these processes (Figure 1F). Next, we further evaluated the applicability of in situ ChIP to non-histone proteins, such as transcription factors (TFs). NKX2-5 as a master cardiac TF is essential for heart development. To showcase TF in situ ChIP in intact tissues, we performed it with a single E9.25 heart each experiment. Compared to NKX2-5 enrichment by standard ChIP-seq of pooling ten hearts for one single experiment, in situ ChIP clearly yielded better signal to noise as showed by track view of two genomic loci and signals at the proximal promoter regions (Supplementary Figure 2A,B). Together, in situ ChIP using low-input materials provided an effective, cost-efficient and easily adapted manner for genome wide profiling of both histone marks and chromatin associated proteins, importantly, which can be finished within one day.

We extended this strategy to assay single cells through the combinatorial indexing (CoBATCH). Permeabilized cells were incubated with antibodies before distribution into one 96-well plate with 200-2,000 cells per well, which were subjected to PAT with different combination of T5 and T7 barcodes for indexing. After the transposition and tagmentation, cells were collected and re-distributed into one or multiple second 96-well plates with 20-25 cells per well, in which different PCR index primers (i5 and i7) were used for the library preparation in the same tube (Figure 2A). In a pilot test, we mixed equal number of mouse ESCs and human HEK293T cells, sequenced to the saturation as determined by the PCR duplicate rate (~80% of unique non-duplicated reads). Given each barcode combination corresponds to either a mouse or human cell, its reads should be mapped predominantly to either the mouse or human genome. We found the collision rate of ~7% (Figure 2B), suggesting a successful de-barcoding of single cells. Using one second 96-well plate for the secondary barcode introduction, we performed single-cell CoBATCH with H3K27ac, and retained 2,161 and 2,388 single cells for further analysis using the conservative cutoff of 3,000 reads per cell. Global analysis displayed a good correlation (Spearman correlation coefficient, 0.90) between native and fixed single-cell experiments (Figure 2C). The aggregate single cells exhibited similar profiling to those generated through standard bulk ChIP-seq in mouse ESCs (Figure 2D). We obtained an average 9,247 unique non-duplicated reads per cell (Figure 2E), close to the yield in recently developed scATAC-seq^4^. Consistent to the lower background as shown by few reads falling in non-peak regions (Figure 2D), the high precision in our data was verified as calculated by fraction of reads in peaks (FRiP) (Figure 2F).

**Figure 2.**
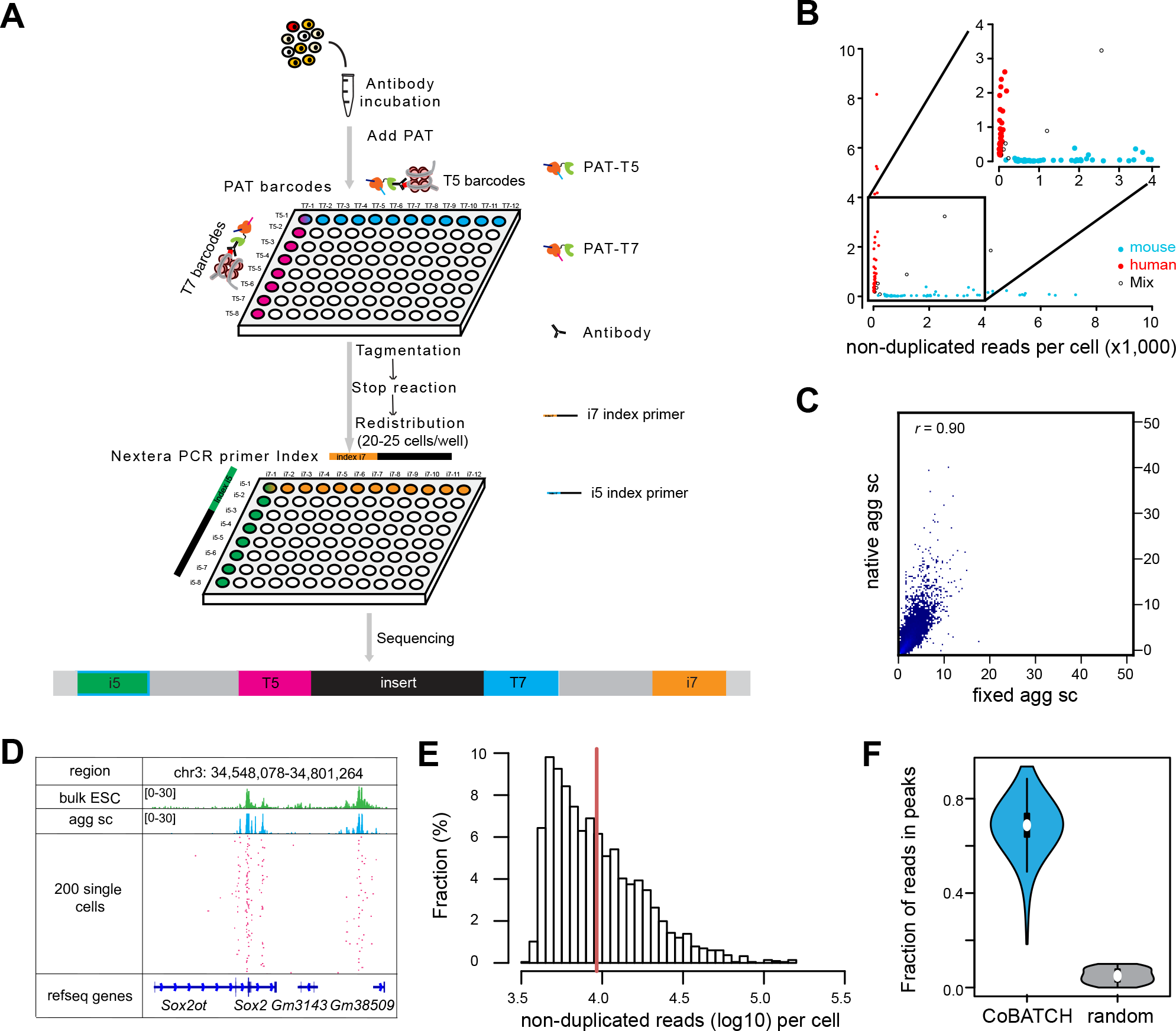
High-throughput CoBATCH prolifing of histone modiciations in single cells under both native and fixed conditions. (A) Schematic of the CoBATCH workflow. (B) Scatter plot showing low collision rate for each unique barcode combination. Mouse ESCs and HEK293T cells were mixed at 1:1. We randomly selected 100 single cells for human-mouse species mix test. (C) Comparison of native and fixed samples using peak regions of aggeragete single cells from H3K27ac CoBATCH data in mouse ESCs. Spearman correlation coefficient was calculated using peak regions. agg sc, aggregate single cells. (D) IGV Track view of H3K27ac signals from both bulk ChIP and CoBATCH data at the representative locus. Aggregated 2,161 single cells and selected 200 single-cell profilings were used for visualization. Bulk H3K27ac ChIP data was from ENCODE. (E) Distribution of non-duplicated reads of 2,161 single cells in mouse ESCs after removal of potential barcodes with less than 3,000 reads.Red line indicates the average non-duplicated reads (9,247 reads per cell) of 2,161 single cells. (F) Violin plot showing the fraction of reads in peak (FRiP) between H3K27ac CoBATCH data in mouse ESCs and simulated random profilings.

We applied single-cell CoBATCH to study epigenetic heterogeneity of a given cell type in tissues. Endothelial cells (ECs) form the vascular system of mammalian bodies, involved in circulation, hematopoiesis, immune, mechanical sensing and organ morphogenesis. The heterogeneity of endothelial cells (ECs) has been studied through the technique of single cell RNA-seq^17^. To obtain a comprehensive epigenetic landscape during ECs differentiation and fate transition for understanding of developmental origins in multiple organs and regulatory correlations with corresponding germ layers, we performed single-cell CoBATCH for H3K27ac in Cdh5 traced EC lineages in 10 organs of E16.5 mouse embryos (Cdh5^CreER^∷Rosa26^tdTomato/+)^ (seen also in Methods). The tdTomato positive cells from different organs were sorted by FACS and fixed before processing (Figure 3A). We obtained an average of ~10,100 unique non-duplicated reads per cell (Supplementary Table 4). Expectedly, both the aggregate single cells and individual single cells from 10 organs exhibited strongest H3K27ac signals at the *Cdh5* enhancer (Figure 3B). We further evaluated CoBATCH data on the tissue-enriched enhancer regions by inspection of aggregate single cells across tissues. Evidently, robust H3K27ac enrichment was produced around the enhancer regions of TFs reflecting tissue developmental origins, *Foxf2* for brain^18^, *Gata4* for heart^19^, *Hoxa11* for kidney^20^, *Foxf1* for lung^21^ and *Cdx2* for small intestine^22^ (Figure 3C). Hierarchical clustering of aggregate single cells showed that limb muscle, skin and small intestine were significantly separated from other tissues. Interestingly, although ECs from right forebrain and hindbrain were expectedly clustered together, ECs from left forebrain were surprisingly far separated and displayed higher similarity to those from liver and lung (Figure 3D). To further examine H3K27ac CoBATCH profiling for the heterogeneities of developmental histories and cell fate transition, we projected all single ECs onto two dimensions by multidimensional scaling (MDS), and found that ECs from heart and kidney were merged together, consistent with the result in Figure 3D, potentially due to their similar epigenetic characteristics of mesodermal derivatives. Altogether, we demonstrated that single-cell CoBATCH in ECs enables mapping of a comprehensive epigenetic landscape for deciphering the intrinsic epigenetic memory reflecting the cell fate dynamics and plasticity of ECs from different mouse organs.

**Figure 3.**
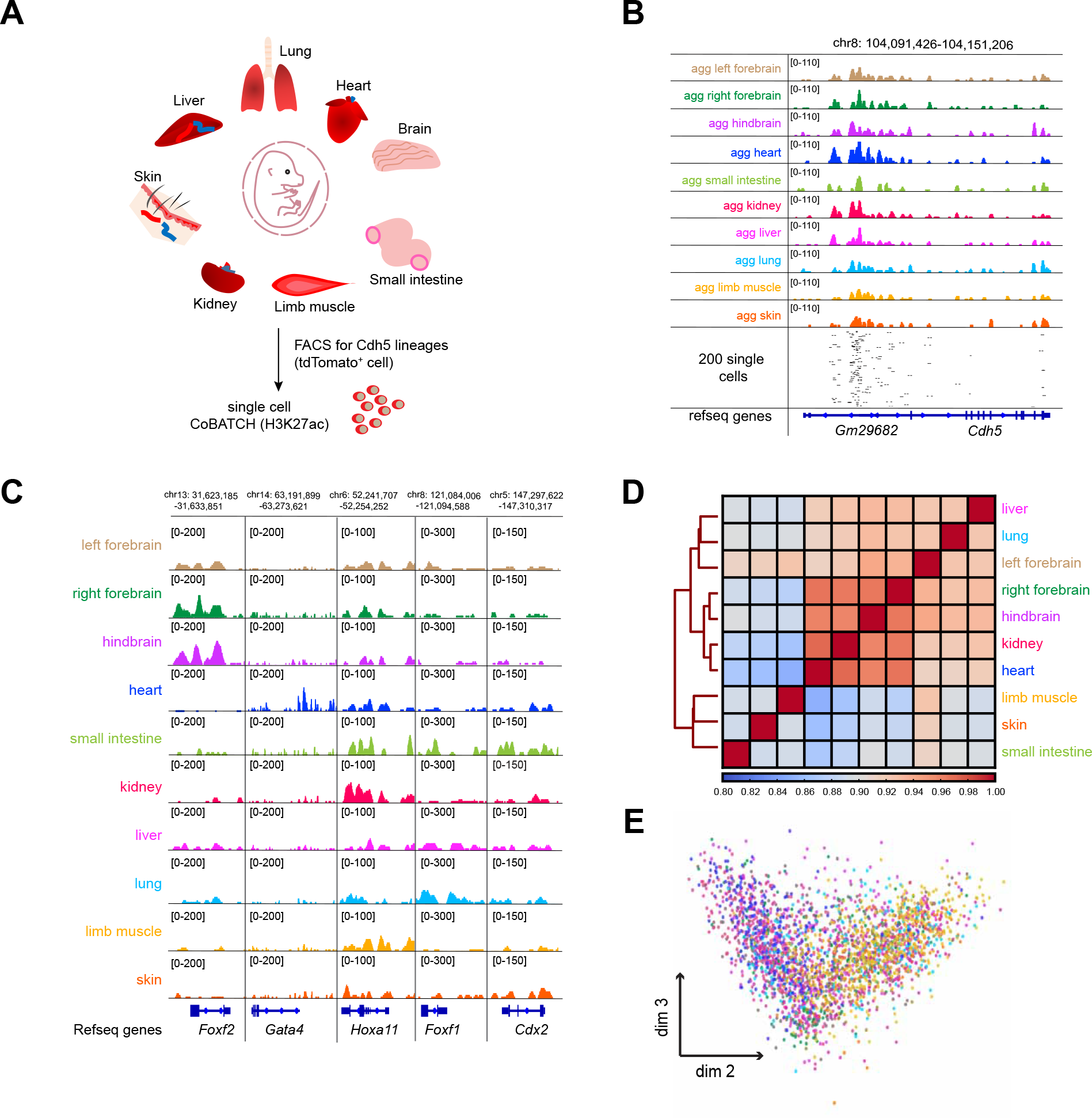
Single cell CoBATCH reveals epigenetic heterogeneity of endothelial cells from 10 mouse embryo organs. (A) Schematic of the single cell H3K27ac CoBATCH experimental design. Endothelial cell (EC) lineages traced by Cdh5^CreER^∷Rosa26^tdTomato/+^ reporter from 10 organs, including heart, liver, lung, forebrain (left), forebrain (right), hindbrain, kidney, skin, limb muscle and small intestine of E16.5 mouse embryos. (B) Track view showing the aggregate profiling of H3K27ac CoBATCH data in each mouse organ at the EC marker gene locus. Single cell H3K27ac profiling was visualized using 200 selected single cells. (C) Track view showing the aggragate H3K27ac CoBATCH data of 10 mouse organs at the organ-specific marker gene loci. (D) Hierarchical clustering of the aggregate single cell profilings of 10 mouse organs by using the genome-wide H3K27ac signals in non-overlapping 5 kb bin windows. (E) Multidimensional scaling (MDS) plot of H3K27ac signals of a total of 2,758 single ECs across 10 mouse organs. We binned the genome by 5 kb windows and calculated the counts of each single cell. The count matrix was transformed into a binary matrix as input for dimension reduction.

In sum, we present an easy-to-use and robust strategy for epigenomic profiling in both low input cells (in situ ChIP) and high-throughput single cells (CoBATCH) in cultured cells and intact tissues for genomic features of TF binding and diverse histone marks. The throughput in CoBATCH can be easily scaled up to ~20,000 single cells by simply redistributing cells into ten 96-well or three 384-well PCR plates during the secondary barcode introduction step. Importantly, our method yields ~10,000 reads per cells with a high signal to noise ratio, requiring a markedly reduced sequencing depth as an advantage of cost-effectiveness. We demonstrated that CoBATCH can be applied to dissect epigenetic heterogeneities and infer developmental origins of a given cell population. Expectedly, this method provides an opportunity to understanding of gene regulation in physiological and pathological processes. We also envision that future improvement toward simultaneous measurement of distinct histone marks or chromatin-binding proteins is achievable through optimizing barcoding strategy. Although high throughput in single cell genomics is in general pursued in most circumstances, the ability to assay small samples as few as tens of cells at single-cell level is equally important, e.g. mapping epigenomic landscape during early mammalian embryo pre-implantation.

## DATA AVAILABILITY

All sequencing data have been deposited to GEO and will be available upon acceptance.

## ACKNOWLEDGMENTS

We thank all members of the He lab for critical comments on this manuscript. A.H. was supported by grants from the National Basic Research Program of China (2017YFA0103402), the National Natural Science Foundation of China (31571487 and 31771607), the Peking-Tsinghua Center for Life Sciences, and the 1000 Youth Talents Program of China.

## AUTHOR CONTRIBUTIONS

A.H. conceived and designed the study. Q.W., S.A and X.Y. designed and performed all experiments. H.X. and Y.L. performed the computational analyses. J.Z. assisted with the experiments. H.X., S.A., Y.L., Q.W. and A.H. wrote the paper with input from all other authors. All participated in data discussion and interpretation.

## METHODS

### ESC culture and preparation for CoBATCH

Wild-type V6.5 murine embryonic stem cells (ESCs) were cultured on 0.1% gelatin-coated plates in ESC DMEM culture medium containing 15% fetal bovine serum (Sigma), 1% Penicillin/Streptomycin (Hyclone), 1% Glutamax (GIBCO), 0.1 mM 2-mercaptoethanol (Sigma), 1% MEM nonessential amino acids (Cellgro), 1% nucleoside (Millipore), and 1,000 U/ml recombinant leukemia inhibitory factor (LIF) (Millipore). For native CoBATCH experiments, cells were dissociated with 0.1% trypsin for 5 min at 37°C and neutralized by FBS. For fixed CoBATCH experiments, cells were dissociated with 0.1% trypsin for 5 min at 37°C and fixed in 1% formaldehyde (Sigma) at room temperature (RT) for 3 min, followed by quenching with 125 mM Glycine (AMRESCO). Fixed cells were washed by 1 ml cold PBS three times and the liquid was removed after centrifugation at 600 g for 3 min at 4°C. Cell pellets can be stored at −80°C or be immediately sorted by FACS for groups with various cell numbers.

### Single cell preparation of endothelial cells from ten mouse E16.5 organs for CoBATCH

Pregnant *VE*^*creERT2*^*∷Rosa26*^*tdTomato/tdTomato*^ mice^*23*^ were sacrificed at E16.5. Organs including heart, liver, lung, left forebrain, right forebrain, hindbrain, kidney, skin, limb muscle and small intestines were micro-dissected and minced into 1~3-mm pieces on ice with sterilized scissors in 1.5 ml tubes containing 1 ml HBSS (with 10 mM sodium butyrate). Tissues were collected by centrifugation at 20 g for 3 min at 4°C and washed once with 1 ml cold HBSS (with 10 mM sodium butyrate) to remove excessive blood. Next, tissues from ten organs were individually digested with 1 ml digestion buffer (Supplementary Table 1 for the specific digestion buffer recipes for different organs) ^24 1^ at 37 °C for 10-25 min in the Eppendorf ThermoMixer with 600 rpm. To aid in digestion, 10 U/ml DNase I (TaKaRa 2270A) was added to the reaction after 3-min digestion and gently pipetting was repeated every 5-6 min digestion. The reaction was neutralized by adding 100 μl %FBS/PBS (Sigma) and cells were collected by centrifugation at 500 g for 3 min at 4°C. To get rid of blood cell, cells were suspended with 1 ml of 1x RBC (erythrocyte lysis buffer) and incubated at RT for 10 min. RBC buffer was removed by centrifugation at 500 g for 3 min at 4°C and the cells were then re-suspended with 1 ml RT pre-warmed HBSS (10 mM sodium butyrate). The dissociated cells were cross-linked with 1% formaldehyde at RT for 3 min and quenched by addition of 55 μl 2.5 M Glycine followed by incubated at RT for 5 min. Samples were quickly washed 3 times with 1 ml cold HBSS. Cells were suspended with 300 μl 1% FBS /HBSS (with 10 mM sodium butyrate) and subjected to FACS (Beckman Coulter MoFlo XDP) sorting for tdTomato positive cells. The positive cells were collected by centrifugation and can be stored at −80°C for later use.

### Cloning of hyperactive protein A-Tn5 into pET28 vector

The protein A coding sequence was amplified from pBS1479 (a gift from Qing Li lab, Peking University) and Tn5 coding sequence was amplified from pTXB1-Tn5 (addgene #60240). Protein A and Tn5 sequences were cloned into pET28a to generate the His-pA-Tn5 (PAT) expression vector.

### Induction of PAT

His-pA-Tn5 (PAT) expression vector was transformed into Bl21 (DE3) chemically competent cells following the standard protocol. One single clone was inoculated into 100 ml LB medium (containing 50 μg/ml kanamycin) in a 500 ml conical beaker and grown at 37°C overnight. 10 ml of over-night grown cells were transferred into 1 L LB medium (containing 50 μg/ml kanamycin) in a 3 L conical beaker and the culture was further incubated at 37°C and 220 rpm for 3-4 h or until the O.D. of up to ~0.8. Bacteria were chilled on ice for 30 min with occasionally shaking. Fresh IPTG was added to the culture to the final concentration of 0.2 mM to induce PAT expression^25^. The culture was incubated at 23°C, 80 rpm for 5 h for induction. Bacteria were harvested by centrifugation with Beckman JXN-26 (JA-10) at 5,000 rpm, and 4°C for 5 min. Cells pellets were washed once by pre-chilled PBS and can be directly used for PAT purification or be stored at −80°C for later use.

### Purification of PAT

Frozen pellets (harvested from 1 L culture) were resuspended thoroughly by 20 ml HXG buffer (20 mM HEPES-KOH, pH 7.2, 0.8 M NaCl, 10% glycerol, 0.2% Triton X-100) supplemented with 1 X EDTA-free protease inhibitor cocktails (Roche) and 1 mM PMSF (AMRESCO). Lysates were sonicated at 8 s ON, 16 s OFF, 20% amplitude, total 5 min (Sonics) in the ice-water mixture. Supernatants were collected at 10,000 rpm, 4°C for 30 min, and 0.1 ml 10% PEI was added (Sigma P3143) dropwise to precipitate bacterial DNA. The PEI-precipitated fraction was removed by centrifugation at 10,000 rpm, 4°C for 10 min and supernatants were collected. To equilibrate Ni-NTA column (QIAGEN), the equilibrium buffer (HXG buffer + 10 mM imidazole) was added to 2 ml Ni-NTA column at a flow rate of 1 ml/min. Cleared lysates were loaded onto the equilibrated Ni-NTA column at a flow rate of 0.7 ml/min. After loading the sample, the Ni-NTA column was washed with 100 ml wash buffer (HXG buffer + 20 mM imidazole) at a flow rate of 1 ml/min. Finally, PAT was eluted from the Ni-NTA column with 5 ml elution buffer (HXG buffer+300 mM imidazole) at a flow rate of 1 ml/min. To remove imidazole, the flow-through fraction was dialyzed in 1 L 2X dialysis buffer (100 HEPES-KOH at pH7.2, 0.2 M NaCl, 0.2 mM EDTA, 2 mM DTT, 0.2% Triton X-100, 20% glycerol) at 4°C overnight. PAT was concentrated by centrifugation using a 30 kDa cutoff ultracentrifuge column (Millipore) and supplied with 1X volume of absolute glycerol (Sigma) before storing at −20°C.

### Quality control for the assembled PAT transposome

The quantity of purified PAT transposase was visualized by Coomassie bright blue staining after proteins were resolved on 7.5% SDS-PAGE gel. Assembly of PAT transposome was described previously ^26 25^. Briefly, all oligonucleotides (Supplementary Table 2) were dissolved in TE buffer (5 mM Tris pH 8.0, 5 mM NaCl, 0.25 mM EDTA) to make 200 μM stock solution. To anneal adaptors, each T5 or T7 oligos were mixed with equal volume of MErev oligo, placed at a thermal cycler for 5 min at 95°C followed by programmed temperature decrease at 0.1°C /s to 75°C, and left in 75°C water to naturally cool down to RT. The 37.5 μM barcoded PAT transposome complex was obtained by adding 37.5 μM annealed adaptors, 37.5 μM transposase PAT and storage buffer (50 mM Hepes pH7.2, 100 mM NaCl, 0.1 mM EDTA, 1 mM DTT, 0.1% Triton X-100, 60% glycerol) and incubating at 25°C for 60 min. The assembled PAT transposome are stable and can be stored −20°C for about half a year.

The activity of assembled PAT complex was examined by tagmentation of mouse genomic DNA as previously described ^26 25^. Briefly, 1 μl 37.5 μM barcoded PAT complex was mixed with 2.5 μl 200 ng/μl mouse genomic DNA, 10 μl ddH_2_O and 4 μl TAPS-MgCl_2_-DMF (50 mM TAPS-KOH pH 8.3, 25 mM MgCl_2_, 50% DMF). Transposition reaction was set up by incubation at 55°C 10 min, add the reaction was stopped by adding 2 μl Stopping buffer (250 mM EDTA, 0.2% SDS) followed by incubation for another 5 min. The fragmented DNA was resolved on 1.5% agarose gel for examination of size distribution. The assembled PA-Tn5 complex yielding fragments with average size no more than 500 bp passed quality control and will be used for the following experiments.

### Low-input cell in situ ChIP

Fresh cells for low-input In situ ChIP can be processed in a single tube within one day. Adequate ESC cells were harvested and washed twice with Wash Buffer (20 mM HEPES pH 7.5, 150 mM NaCl, 0.5 mM Spermidine, 1X cocktail (Roche), 1 mM PMSF (AMRESCO) and 10 mM sodium butyrate). Concanavalin A coated magnetic beads (Con-A beads) (Bangs Laboratories) were prepared as described. Briefly, adequate Con-A beads were washed with 1 ml Binding Buffer (400 μl 1M HEPES pH 7.9, 200 μl 1M KCl, 20 μl 1M CaCl2, 20 μl 1M MnCl2, and the final volume to 20 ml with ddH_2_O) 1-2 times, resuspended with adequate Binding Buffer and applied to the cell mixture. The cells and Con-A beads mixture were incubated at RT for 10 min and collected by the magnet stand. Next, cells were resuspended with 100 μl Antibody Buffer (mix 4 μl 0.5 M EDTA with 1 ml 0.01% Digitonin-Wash Buffer (Dig-Wash Buffer)) supplemented with cocktail, PMSF, 10 mM sodium butyrate and 0.1% Triton X-100) containing 0.5 μl antibody by gently flicking (Supplementary Table 3). The cell and antibody mixture were incubated at 4°C for 1-2 h. The liquid was removed from the caps and the sides of the tubes with a quick pulse on a microcentrifuge (<100 g, 22 °C, 1 s). Cells were washed twice with 180 μl Dig-Wash Buffer (with 0.1% TX-100) and finally suspended with 100 μl Dig-Wash Buffer (with 0.1% TX-100, cocktail, PMSF and 10 mM sodium butyrate) containing 3 μg/ml PAT-MEAB. The cells were incubated at 4°C for 1 h and washed with 180 μl Dig-Wash Buffer with 0.1% Triton X-100 three times for 5 min each to get rid of free PAT-MEA/B. The reaction was activated by suspending the cells with 25 μl cold Reaction Buffer (10 mM TAPS-NaOH pH 8.3, 5 mM MgCl2, 10% DMF and supplemented with cocktail, PMSF 10 mM sodium butyrate) by gently flicking and the reaction was incubated at 25°C for 1 h in a PCR cycler. The reaction was gently mixed once after 30-min incubation. To stop the reaction, 10 μl 40 mM EDTA was added followed by further incubation at RT for 15 min after mixing well. Samples were washed once with 1% BSA/PBS with 1mM EDTA and collected by a magnetic stand. Samples were resuspended with 5 μl Lysis Buffer containing 10 mM Tris-HCl pH 8.5, 0.05 % SDS and 0.1 mg/ml Proteinase K (AMRESCO). Samples were incubated at 65°C for 1 h to release DNA fragments and then incubated at 85°C for 15 min to deactivate Proteinase K. 1 μl 1.8% Triton X-100 was added before incubation at 37°C 1 h to quench SDS in the reaction. DNA in the reaction can be directly used, without need of purification, for library preparation in the same tube as blows:

### Nextera library preparation and sequencing strategy of low-input in situ ChIP

PCR amplification was performed by addition of 1 μl 25 μM i5 index primer (5′- AATGATACGGCGACCACCGAGATCTACAC[i5]TCGTCGGCAGCGTC-3′), 1 μl 25 μM i7 index primer (5′- CAAGCAGAAGACGGCATACGAGAT[i7]GTCTCGTGGGCTCGG-3′) (Supplementary Table 2), 25 μl 2X KAPA master mix and 17 μl ddH_2_O to the 6 μl CoBATCH cell lysis and the reaction was set up by incubation at 72°C for 5 min, 98°C for 45 s, 10-18 cycles of (98°C for 15 s, 63 °C for 30 s, 72°C for 1 min), and final 72 °C extension for 5 min. After PCR, the library was purified with 1 X AMPure XP beads once. Size selection was carried out by first 0.5X AMPure XP beads to remove >1kb fragments, and second 0.5X AMPure XP beads to the supernatant to obtain 200-1000 bp fragments for sequencing. The libraries were sequenced with paired-end 150-bp reads on Hiseq X-ten or Novaseq 6000 platform (Illumina).

### Fixed single-cell CoBATCH

Fixed cells were resuspended with 200 μl Hypotonic Buffer (20 mM HEPES pH 7.9, 10 mM KCl, 10% Glycerol, 0.2% NP40, 0.3% SDS) supplemented with adequate proteinase inhibitor cocktail, PMSF and 10 mM sodium butyrate. Chromatin was opened at 62°C for 10 min in an Eppendorf ThermoMixer at 600 rpm, quenched by adding 20 μl 20% Triton X-100 and incubating at 37°C for 1 h in an Eppendorf ThermoMixer at 600 rpm. Cells were washed twice with 1 ml Wash Buffer (1 ml 1 M HEPES pH 7.5, 1.5 ml 5 M NaCl, 12.5 μl 2 M spermidine, 10 mM and sodium butyrate, and the final volume to 50 ml with ddH_2_O) and suspended with 1 ml Wash Buffer containing 10~50 μl activated Con-A beads. The cells-ConA beads mixture was incubated at RT for 10 min for binding before collected by a magnet stand. Cells were resuspended in 100 μl Antibody buffer with 0.5 μl antibody by gently vortexing (Supplementary Table 3) and incubated at 4°C for 2~4 h. After antibody binding, the cells were washed twice with 180 μl Dig-Wash Buffer and resuspended with 1% BSA/PBS. The 96-well plates were prepared for PAT-barcodes complex binding: the 96-well plates were washed with 1% BSA/PBS once followed by PBS wash. 100 μl Dig-Wash Buffer containing individual combinatorial barcoded 3 μg/ml PAT-T5 and 3 μg/ml PAT-T7 (Supplementary Table 2) supplemented with proteinase inhibitors and 10 mM sodium butyrate was added to each well of the pre-washed 96-well plates. Cells in 1% BSA/PBS was FACS sorted for 2000 cells into each well. The 96-well plates were incubated at 4°C for 1 h for PAT-barcode complex binding followed by washing with 180 μl Dig-Wash Buffer (with 0.1% Triton X-100) twice. Cells were resuspended with 10 μl cold Reaction Buffer (10 mM TAPS-NaOH pH 8.3, 5 mM MgCl2, 10% DMF and supplemented with cocktail, PMSF 10 mM sodium butyrate) by gently vortexing. The reaction was activated by incubating the plates at 37°C for 1 h on the PCR cycler. The reaction was stopped by adding 10 μl 40 mM EDTA to each well and mixing well. The plates were further incubated at RT for 15 min and 20 μl cold Sort Buffer (PBS / 2% BSA / 2 mM EDTA) was added to each well. Cells from the 96-well plates were combined to a new tube for DAPI staining (Thermo). Cells were passed through a 30 μm cell strainer to remove cell clumps, and 20-25 cells were sorted into each well of a new 96 well plate which contains 4 μl Lysis Buffer (10 mM Tris-HCl pH 8.5, 0.05 % SDS and 0.1 mg/ml Proteinase K). The 96-well plates were incubated cells at 65°C overnight for 5-6 h for reverse crosslinking and incubated at 85°C for 15 min to deactivate proteinase K. 1 μl 1.8% Triton X-100 was added to each well and the reaction was incubated at 37°C for 1 h to quench SDS. The PCR enrichment was directly performed in the same tube as described above.

### Native single-cell CoBATCH

Native single-cell Co-BATCH was performed as following the fixed CoBATCH protocol, except for the elimination of fixation process and lysis at 65°C for 10 min.

## BIOINFORMATICS

### Low-input in situ ChIP data processing and peak calling

Firstly, we evaluated the quality of in situ ChIP-seq data by FastQC (version 0.11.5). Removing adapters and trimming low-quality bases at the end of reads were performed by cutadapt (version 1.11) with the parameters (-q 20 -O 10 -- trim-n -m 30 --max-n 0.1). Clean paired-end in situ ChIP-seq sequences were then mapped to mouse reference genome mm10 using Bowtie2 (version 2.2.9)^27^ with the parameter, “--dovetail --very-sensitive-local --no-unal --no-mixed --no-discordant -X 2000” for sensitive and fast alignment. The mapped reads with MAPQ greater than 30 were identified as uniquely mapped reads, which were sorted using Samtools (version 1.9) and used for the subsequent analyses. PCR duplicates were removed by Picard (version 2.2.4) (http://broadinstitute.github.io/picard) by default parameters. Only uniquely mapped, non-duplicated reads were used for peak calling. Histone modifications’ peaks were identified from antibody group versus IgG group using MACS2 (verison 2.1.1) (PMID: 18798982) with the parameters (--broad) for low-input in situ ChIP-seq. The peaks were annotated by ChIPseeker (version 1.18.0)^28^.

### Single-cell CoBATCH data processing

Single cell CoBATCH libraries were de-multiplexed by in-house scripts. Single cell libraries were processed to generate unique and non-duplicated libraries as mentioned above. For single cell H3K27ac libraries of endothelial cells from 10 mouse embryo organs, we excluded single cells with less than 3,000 non-duplicated reads. We generated peak files using MACS2 with default settings in aggregated bam files in each organ. FRiP (fraction of reads in peaks) was calculated by the non-duplicated reads overlapping with peaks divided by the total non-duplicated reads in each single cell library. Simulated random profiling for FRiP were plotted together as control in violin plot. We plotted the scatter plot in which each point represents the reads mapped to human genome (y axis) and the reads mapped to mouse genome (x axis) in a single cell library by randomly sampled 100 cells in the mixed library which contained 1:1 human and mouse single cells. We defined a “collision” situation as the fraction of reads mapped to human genome (hg19) was between 0.2 and 0.8^3^. These resulted in H3K27ac profiles of total 3,057 single endothelial cells after filtering as above. The specific number varies in endothelial cells from 10 organs: 351 cells in left forebrain, 287 cells in right forebrain, 305 cells in hindbrain, 215 cells in skin, 433 cells in kidney, 389 cells in liver, 235 cells in lung, 288 cells in heart, 74 cells in intestine, and 480 cells in muscle. We used “multiBamSummary” function in deepTools to count the reads of each single cell in genome-wide 5-kb bins, resulting in a region x cell matrix. Next, we binarized the matrix, and filtered out the lowest 10% of cells in terms of number of bins, which was used as input for dimensionality reduction. Finally, we performed MDS reduction and visualization as showed in Figure 3.

### Visualization of low-input In situ data and single-cell CoBATCH data

For visualization of low-input in situ ChIP-seq data, we used bamCoverage in deepTools (version 2.2.3)^29^ to calculate coverage in continuous 50-bp bins in bam files and generate track files (bigwig format). We set the scale factor to normalize sequencing depth of each library to 10,000,000 reads. For single-cell CoBATCH reads visualization, we randomly selected single-cell bigwig files and uploaded into IGV (version 2.3.59)^30^ together with aggregated and reference tracks.

### Receiver operating characteristic (ROC) curve

To evaluate the low-input in situ ChIP-seq data quality, we performed Receiver Operating Characteristic. We carefully chose ESC H3K4me3 ChIP-seq peaks as gold standard from ENCODE database^31^. These H3K4me3 ChIP-seq peaks within regions of ± 2 kb of gene TSS were defined as standard positive peaks. True positive peaks were defined as low-input in situ ChIP peaks overlapping with standard positive peaks. We defined regions in ± 2 kb of gene TSS which have no overlap with ENCODE H3K4me3 ChIP-seq peak as standard false positive regions. False positive peaks were defined as low-input in situ ChIP peaks not overlapping with standard positive peaks. According to this definition, we calculated true positive rate (TPR) as the number of true positive peaks divided by the number of standard positive peaks and false positive rate (FPR) as the number of false positive peaks divided by the number of standard false positive regions. To plot the ROC curve, we set a series of p-value cutoff for peaks and generated a vector comprising TPR and FPR. For each group of different cell number levels, we plotted ROC curve and calculated Area Under Curve (AUC) in R (version 3.3.3)^11^.

### Correlation analysis of in situ ChIP-seq data

To evaluate correlations between H3K4me3 in situ ChIP-seq replicates performed in each stage of embryo, we calculated the normalized average scores for each replicate in every TSS ± 2 kb region by “multiBigwigSummary BED-file” function in deepTools. The Spearman correlation was calculated between replicates and plotted by “plotCorrelation” function. To evaluate correlations between H3K27ac in situ ChIP-seq replicates performed in each stage of embryo, we calculated the normalized average scores for each replicate in continuous 5-kb windows genome wide by “multiBigwigSummary bins” function in deepTools. The Pearson correlation was calculated between replicates and plotted. To evaluate correlations between H3K27ac native CoBATCH and H3K27ac fixed CoBATCH performed in ESCs, we calculated the normalized average scores for each replicate in H3K27ac peaks downloaded from ENCODE by “multiBigwigSummary BED-file” function in deepTools. The Spearman correlation was calculated between replicates and plotted.

---

**Supplementary Figure 1.**
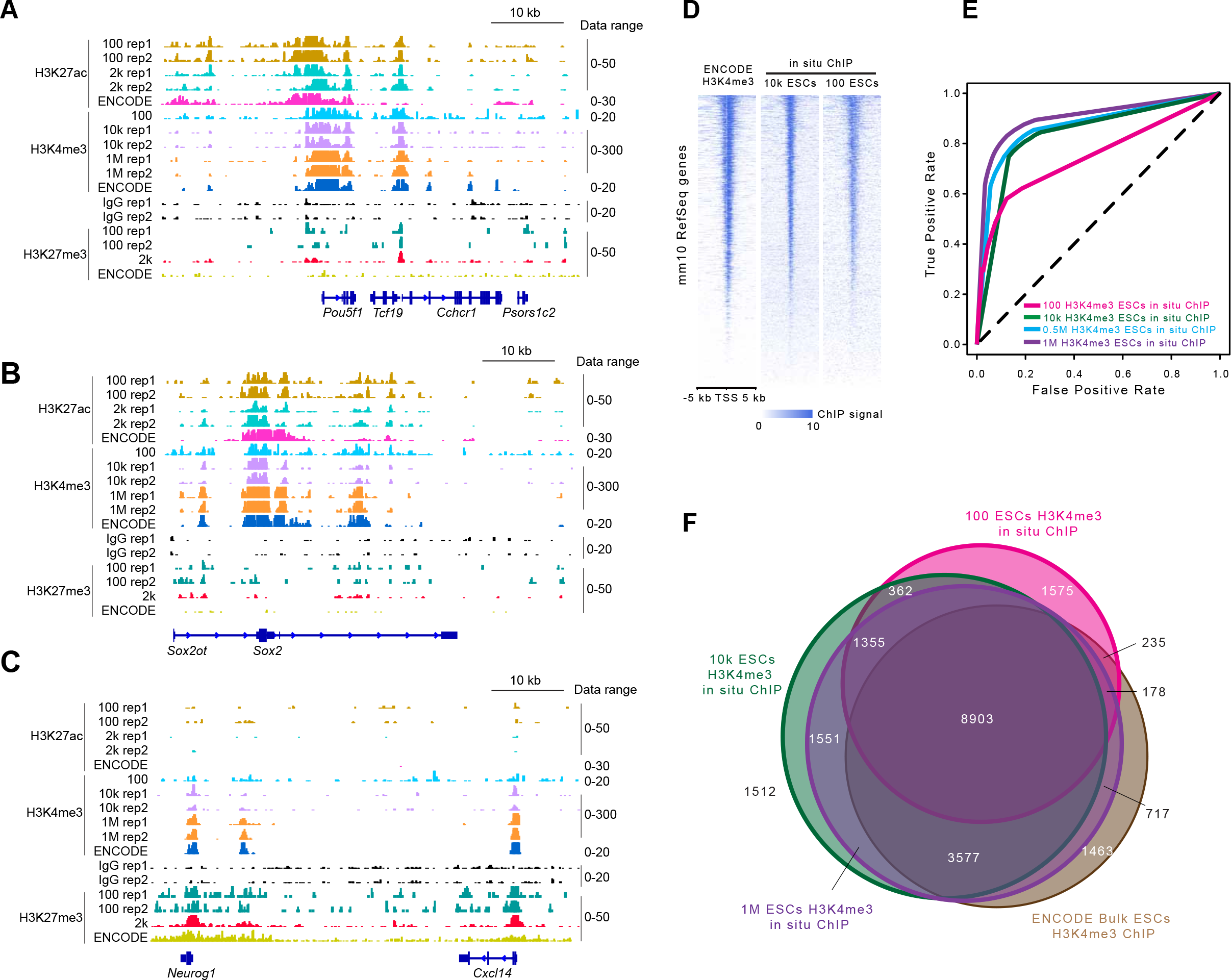
Characterization of low input in situ ChIP of histone marks. **(A, B** and **C)** IGV Track view showing H3K27ac, H3K4me3, and H3K27me3 signals at the representative loci by in situ ChIP with varying cell numbers of ESCs (100, 2k and 10k). Cell numbers are indicated as shown above. (**D**) Heatmaps showing H3K4me3 in situ ChIP signals at gene TSS ± 5 kb regions with indicated cell numbers of ESCs. (**E**) Receiver operating characteristics (ROC) curves showing performance of low-input H3K4me3 in situ ChIP-seq. Gold standard is shared peaks among four ENCODE H3K4me3 ChIP-seq peak datasets. The AUC value of each curve is 0.900 (purple), 0.865 (deepskyblue), 0.835 (green), and 0.748 (meganta) in H3K4me3 in situ ChIP data from 1 millilion (M), 0.5M, 10k, and 100 ESCs, respectively. (**F**) Venn diagram showing peak overlap in groups as in (B). Peaks were called as detailed in the methods. Numbers indicating overlapping peak numbers.

**Supplementary Figure 2.**
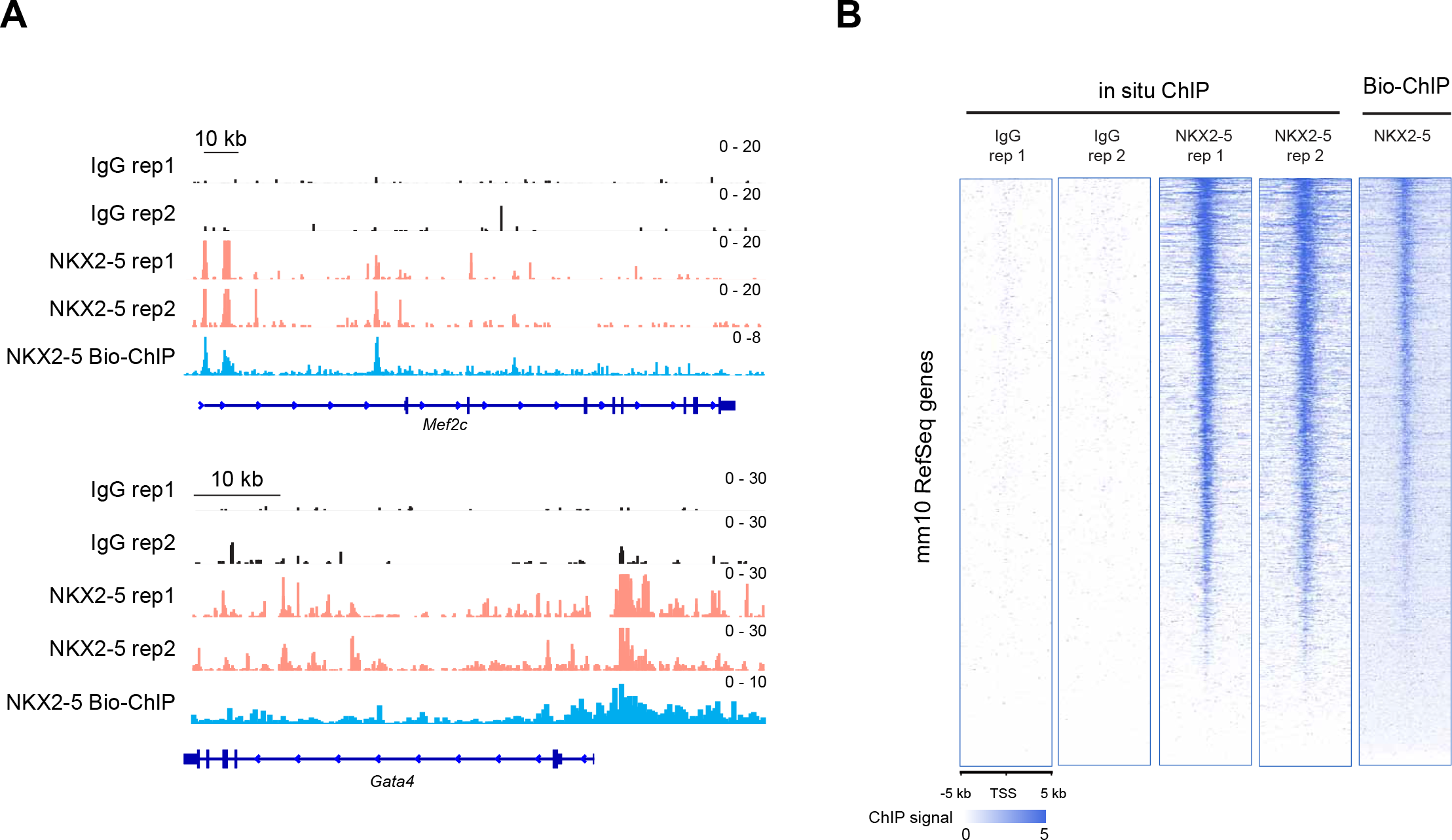
Characterization of NKX2-5 in situ ChIP for embryonic E9.25 heart. **(A)** IGV track view showing NKX2-5 signals at the representative loci by NKX2-5 in situ ChIP of individual embryonic E9.25 hearts (~ 4,000 cells). **(B)** Heatmap showing normalized in situ ChIP signals at gene TSS ± 5 kb regions. Two replicates of IgG in situ ChIP and two replicates of NKX2-5 in situ ChIP were performed in native mouse E9.25 heart and the NKX2-5 Bio-ChIP was performed in fixed E9.25 hearts.

